# Sex classification using long-range temporal dependence of resting-state functional MRI time series

**DOI:** 10.1101/809954

**Authors:** Elvisha Dhamala, Keith W. Jamison, Mert R. Sabuncu, Amy Kuceyeski

## Abstract

A thorough understanding of sex differences, if any, that exist in the brains of healthy individuals is crucial for the study of neurological illnesses that exhibit differences in clinical and behavioural phenotypes between males and females. In this work, we evaluate sex differences in regional temporal dependence of resting-state brain activity using 195 male-female pairs (aged 22-37) from the Human Connectome Project. Male-female pairs are strictly matched for total grey matter volume. We find that males have more persistent long-range temporal dependence than females in regions within temporal, parietal, and occipital cortices. Machine learning algorithms trained on regional temporal dependence measures achieve sex classification accuracies of up to 81%. Regions with the strongest feature importance in the sex classification task included cerebellum, amygdala, frontal cortex, and occipital cortex. Additionally, we find that even after males and females are strictly matched on total grey matter volume, significant regional volumetric sex differences persist in many cortical and subcortical regions. Our results indicate males have larger cerebella, hippocampi, parahippocampi, thalami, caudates, and amygdalae while females have larger cingulates, precunei, frontal cortices, and parietal cortices. Sex classification based on regional volume achieves accuracies of up to 85%; cerebellum, cingulate cortex, and temporal cortex are the most important features. These findings highlight the important role of strict volume matching when studying brain-based sex differences. Differential patterns in regional temporal dependence between males and females identifies a potential neurobiological substrate underlying sex differences in functional brain activation patterns and the behaviours with which they correlate.

## Introduction

The study of sex differences in the brain is one of the most long-standing and debated themes in neuroscience. A compelling reason to investigate sex differences in the brain is that for many neurodevelopmental, neuropsychiatric, and neurodegenerative illnesses, the age of onset, prevalence, and symptomatology varies between the sexes. Furthermore, insight into the etiology of sex differences in the healthy brain provides an important foundation with which to delineate sex-specific pathophysiological mechanisms in different disorders and to guide the development of sex-specific treatment.

Even at rest, the brain generates an ever-changing pattern of activity that can be measured using fMRI. This activity is characterised by long-range temporal dependence such that signal fluctuations at the present time influence signal dynamics up to several minutes in the future. The Hurst Exponent (HE) is a scalar measure of long-range temporal dependence of time series that quantifies the tendency of a time series to either regress to the mean or persistently cluster in a direction [1]. The value of HE can be used to characterise the fMRI signal into three different scenarios: (1) when 0.5 < HE < 1 there exist long-range temporal correlations such that a high value in the time series is likely to be followed by a second high value and this tendency to cluster in one direction relative to the mean is likely to persist in the future; (2) when H=0.5 there exists uncorrelated temporal activity such that the time series is similar to random noise; and, (3) when 0 < HE < 0.5 there exist long-range anticorrelations such that a high value in the time series is likely to be followed by a low value and then another high value, and this tendency to fluctuate between high and low values is likely to persist into the future [1]. Based on these characteristics of the HE, it has been suggested that HE can represent signal complexity of brain activity, with a higher HE corresponding to lower signal complexity [2].

The use of HE to study signal fluctuations in functional MRI (fMRI) in both healthy and clinical populations has recently emerged [1–5]. In recent years, it has been observed that a baseline state of wakefulness in healthy subjects is associated with critical dynamics in the resting state time series and unconscious brain states (e.g. asleep or sedated) exhibit a departure from critical dynamics [1]. This departure corresponds to a reduction in long-range temporal dependence in the frontal lobe, salience network, and thalamus during unconsciousness [1], suggesting that there is a relationship between consciousness and regional temporal dependence in the brain. Another recent study looked at the relationship between long-range temporal dependence and sex in healthy subjects and observed sex differences in frontal, parietal, occipital, and limbic lobes [3]. In order to understand clinical or behavioral implications of HE, we must first fully quantify any baseline sex differences in the temporal dependence of resting-state time series in healthy adults.

In recent years, machine learning techniques have increasingly been used in the analysis of resting-state fMRI data [6]. Supervised methods have been successfully applied to make subject-level predictions in both healthy and clinical populations [6]. Despite the widespread interest on sex differences in the brain, very few studies have focused on sex classification using structural and/or functional information from the healthy brain. Using whole-brain functional connectomes and a partial least squares approach applied to data from the Human Connectome Project (HCP), sex classification has been performed with a maximum accuracy of 79-86% [7]. Another study using a subset of the HCP data compared sex classification accuracies of 436 different models, each one based on a region’s connectivity to the rest of the brain, in an effort to reduce the overall dimensionality. They reported mean accuracies of 60.0% to 68.7%, and a maximum accuracy of 75.1% [8]. While sex classification using functional connectivity profiles has achieved varying levels of accuracy [7, 8], it has not yet been attempted, to the best of our knowledge, using temporal dependence measures of brain activation like HE.

The present study evaluated sex differences in regional long-term temporal dependence of the resting-state fMRI time series in a grey-matter-volume-matched subset of 390 subjects from the HCP [9] dataset. The project’s primary goals were: 1) to quantify regional sex differences in temporal dependence of cortical and sub-cortical grey matter regions and 2) to test if regional temporal dependence information is sufficient to successfully classify subjects based on sex. As a secondary goal, this study also evaluated whether regional volumetric sex differences persisted even after matching subjects on total grey matter volume.

## Results

We present results from a cohort of 390 grey-matter-volume-matched healthy young adults (ages 22-37; 190 males) from the HCP – Young Adult S1200 dataset [9]. Each subject had four complete resting state fMRI scans with 1200 volumes each that were acquired over two sessions and preprocessed [10]. Each of the four fMRI scans was divided into 8 segments of 150 volumes each, and HE was computed using detrended fluctuation analysis [1] on each segment. Voxel-wise HE maps were obtained by averaging HE over all segments from each of the four scans (8 × 4 = 32 segments total) to improve signal-to-noise ratio. Regional HE was obtained by averaging the HE values of each voxel within a given region. Two sample t-tests were used to evaluate regional sex differences in HE and all p-values were corrected for multiple comparisons [11]. A linear support vector machine (SVM) was optimised using 100 iterations of nested grid search with five-fold inner and outer cross validation and fit on a training subset of the total data (80%). Classification performance is calculated based on the final model’s performance on the remaining 20% hold-out test data. This procedure was repeated over 10 permutations to assess robustness to training/test set assignments. To ensure that results were not subjected to biases introduced by a single grey matter parcellation scheme, all analyses were replicated over seven different atlases. An ensemble model that took the average of each of the individual atlas model predictions was also created. Finally, to quantify the effect of sample size and volume-matching on results, HE-based sex classification using the CC400 atlas was also performed: i) for the entire dataset (n=1003) and ii) for 10 randomly selected (not volume-matched) subsets of equal size to the volume-matched sample (n=390). Finally, regional volume was used to perform sex classification to determine whether volume alone, even after strict grey matter volume matching, can still identify sex. Details are given in Materials and Methods and an outline of the workflow can be seen in Figure S1.

### Sex Differences in HE and Volume

Significant sex differences (p-corrected<0.05, correction performed over 1114 regions in all atlases) in HE was observed in 80 out of 1114 total cortical and subcortical grey matter regions across all seven atlases. In all regions found to be significantly different between the sexes, males exhibited a higher HE than females. Group average HE values for the CC400 atlas are shown in Figure 1, with males in panel A and females in panel B. The t-statistics for regions exhibiting significant sex differences in HE for the CC400 atlas are shown in Figure 2a, and for all other atlases in Figure S2. For the CC400 atlas, HE was significantly higher in males than females in 28 regions located in the temporal, parietal, and occipital cortices. The other atlases demonstrate similar regional patterns of significant sex differences.

**Figure 1:**
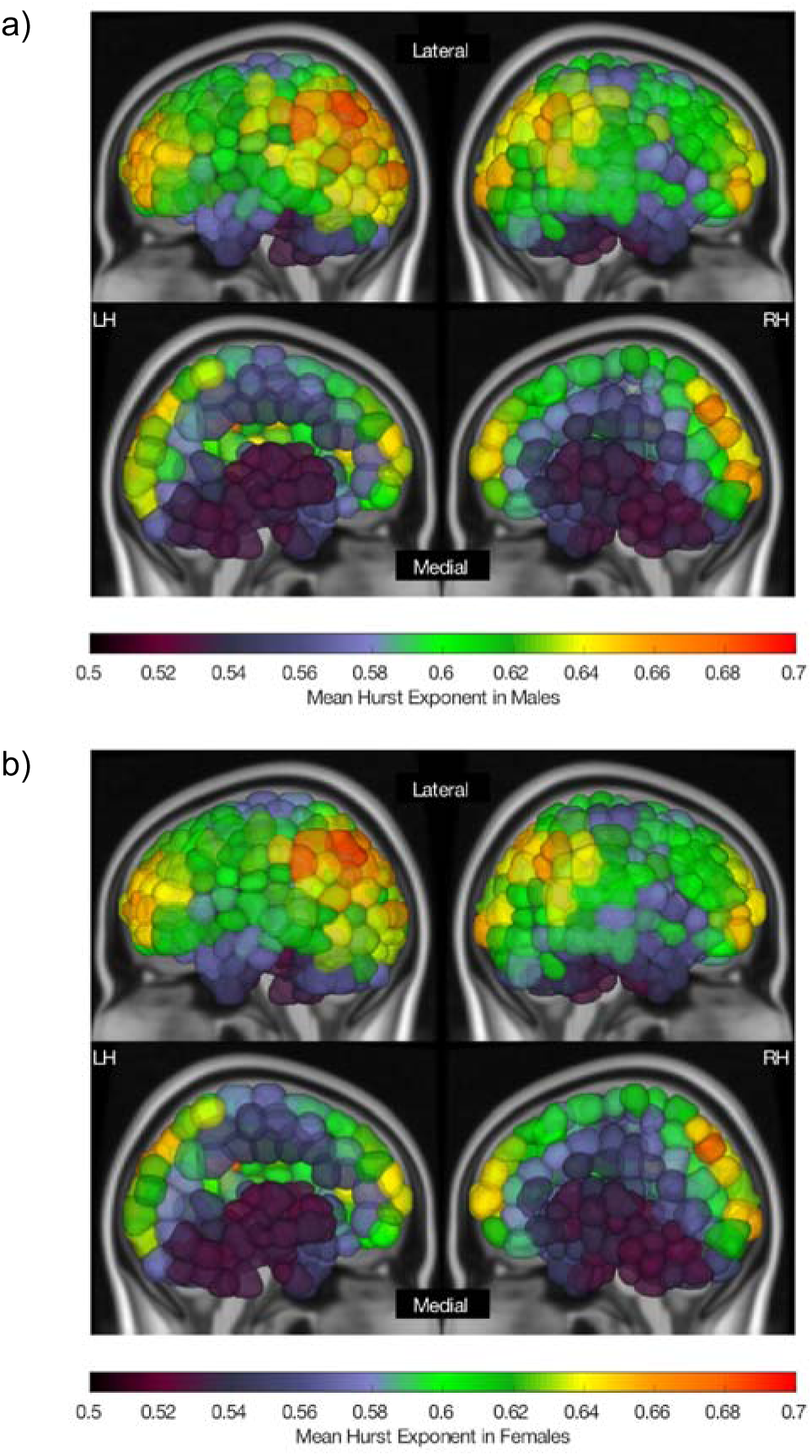
Regional mean HE in males (a) and females (b) computed on the CC400 atlas. Lateral (top) and medial (bottom) sides of the left (LH) and right (RH) hemispheres are shown.

**Figure 2:**
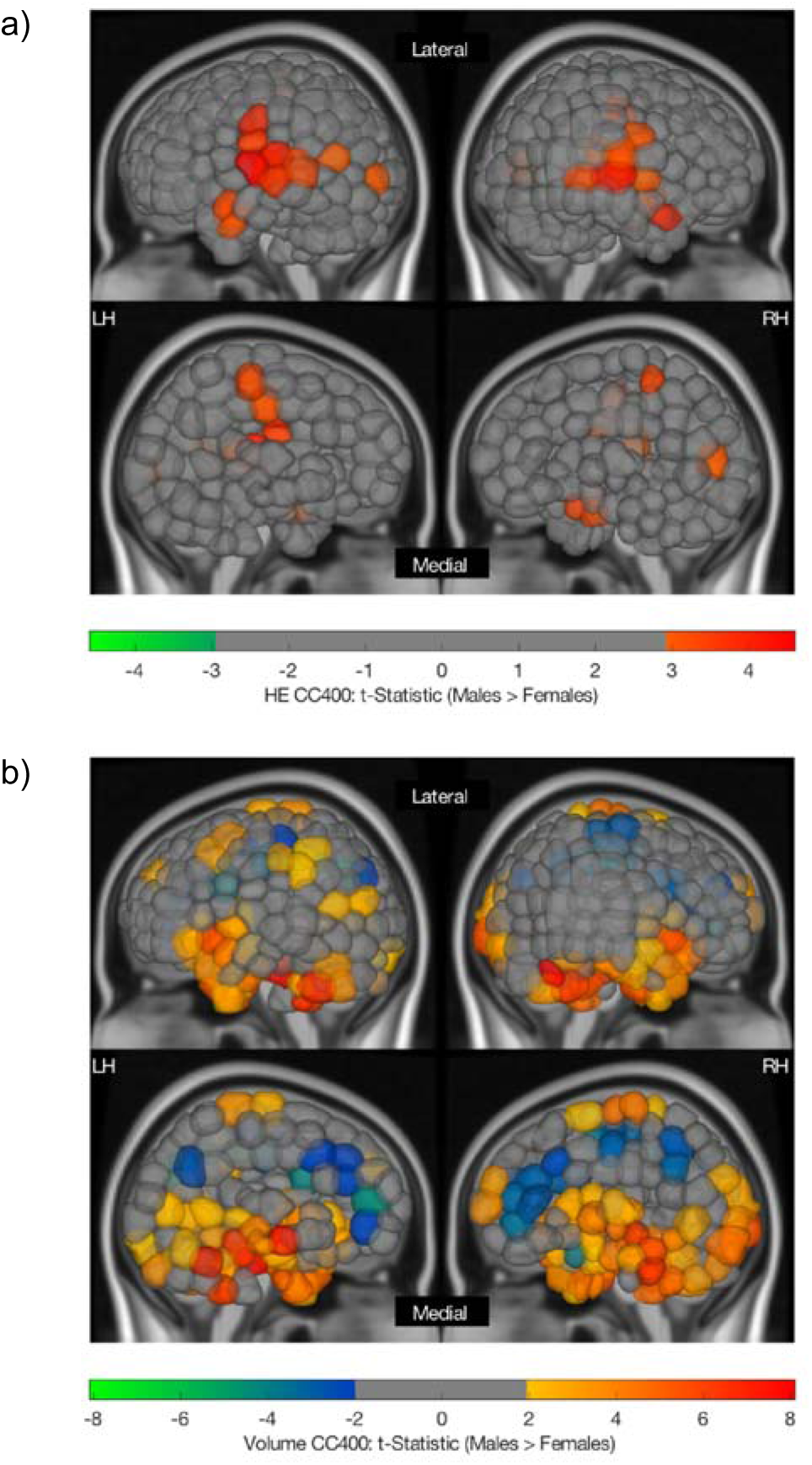
Region-wise sex differences in HE (a) and volume (b) computed on the CC400 atlas. Lateral (top) and medial (bottom) sides of the left (LH) and right (RH) hemispheres are shown for both a and b. Regional t-statistics are shown as per the colour scale for all significantly different (p-corrected < 0.05) areas. Non-significant areas are shown in grey. A positive t-statistic indicates that males have a higher mean value in that region than females.

Significant sex differences (p-corrected<0.05, correction performed over 1114 regions in all atlases) in regional volume was observed in 380 out of 1114 total cortical and subcortical grey matter regions across all seven atlases. Females exhibited larger volumes in the bilateral cingulate cortices, pre- and post-central gyri, precunei, and frontal gyri, while males exhibited larger volumes in bilateral cerebella, hippocampi and parahippocampi, thalami, caudates, and amygdalae. The t-statistics for regions exhibiting significant sex differences in volume for the CC400 atlas are shown in Figure 2b, and for all other atlases in Figure S3. Again, the other atlases demonstrate similar patterns of volumetric sex differences.

### Sex Classification

Receiver operating characteristic (ROC) curves for all atlases for HE-based sex classification can be seen in Figure 3a, and for volume-based sex classification in Figure 3b. Accuracy and area under the ROC curve (AUC) metrics for all atlases can be found in Table 1 for HE-based sex classification and Table 2 for volume-based sex classification. For HE-based classification, mean accuracies ranged from 73.0% to 81.2%, and mean AUCs ranged from 0.79 to 0.87. The CC400 atlas was the best performing atlas for both HE-based and volume-based sex classification. For HE-based classification, it performed with a mean accuracy of 81.2% (SD=3.5%) and a mean AUC of 0.87 (SD=0.033). For volume-based classification, it performed with a mean accuracy of 85.4% (SD=2.5%) and a mean AUC of 0.92 (SD=0.023). The ensemble model combining prediction probabilities from all atlases performed with a mean accuracy of 81.2% (SD=3.4%) and a mean AUC of 0.86 (SD=0.033). For volume-based classification, mean accuracies ranged from 72.2% to 85.4%, and mean AUCs ranged from 0.78 to 0.92. The ensemble model performed with a mean accuracy of 81.4% (SD=4.8%) and a mean AUC of 0.91 (SD=0.033).

**Figure 3:**
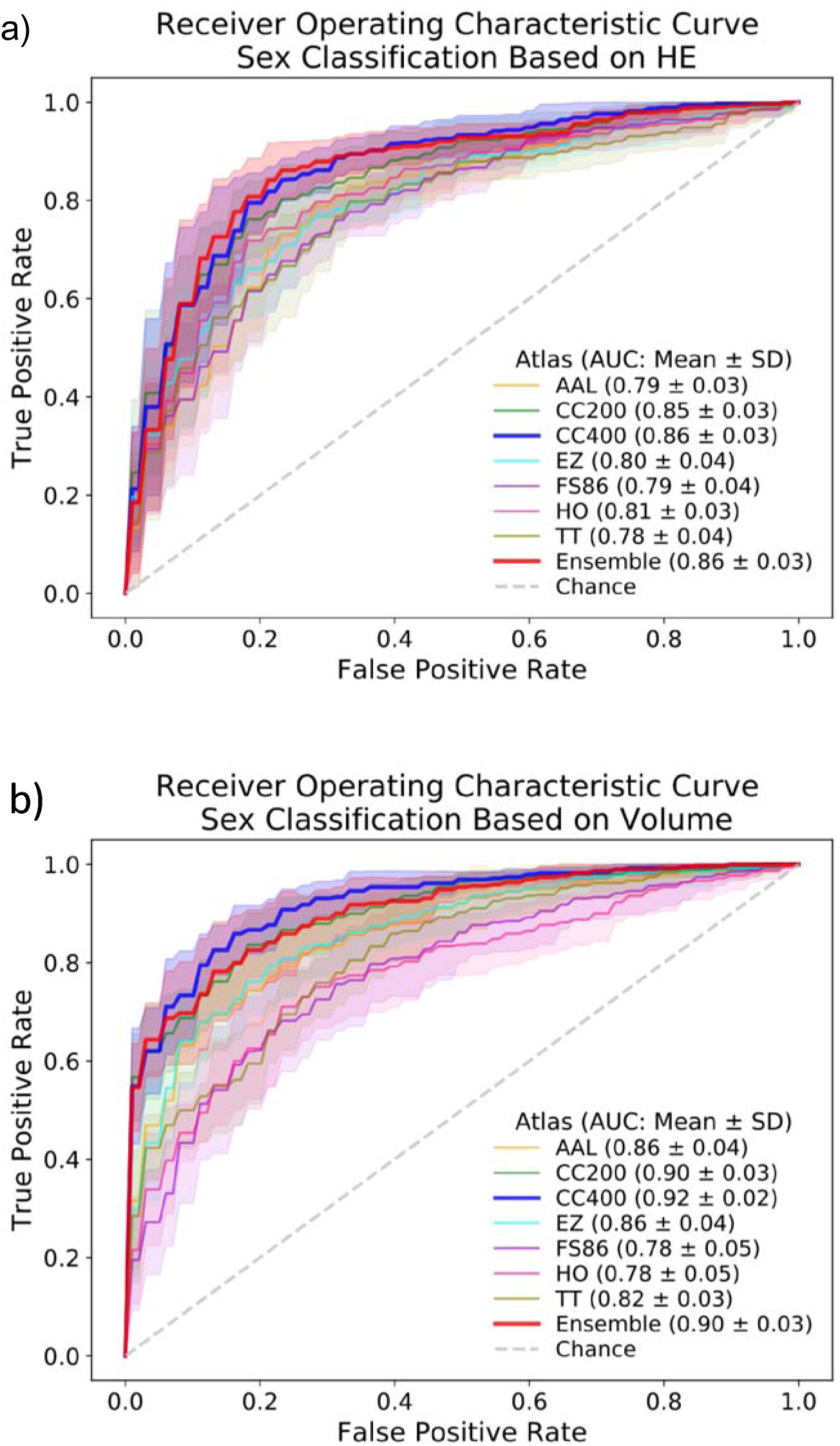
Receiver Operating Characteristic Curve for linear SVM sex classification for all atlases based on HE (a) and volume (b). Mean and standard deviation of the AUC values for each atlas are indicated.

**Table 1:**
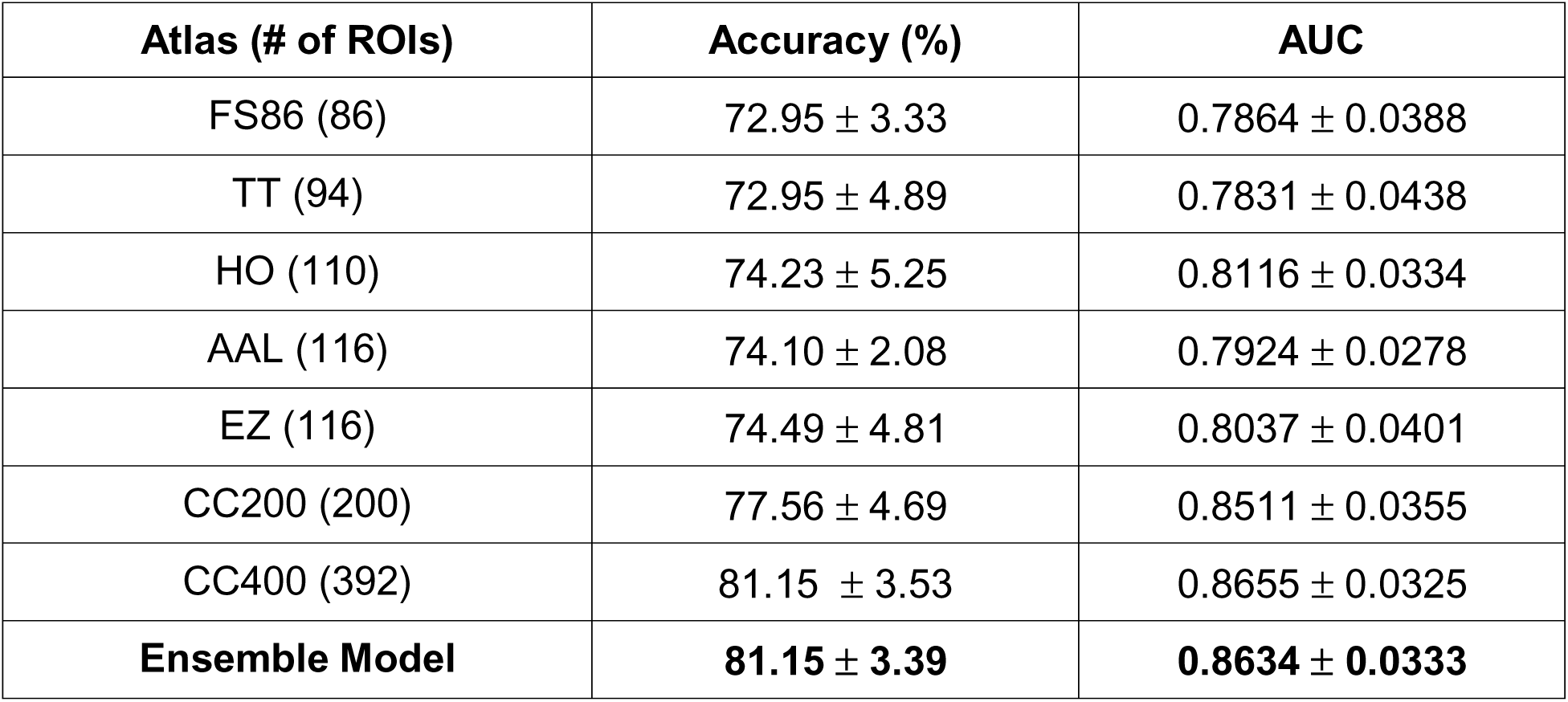
Results from Grey Matter Volume Matched Dataset (n=390). Using HE to predict sex. Ten permutations completed for each atlas. Mean accuracy (balanced) ± standard deviation and mean AUC ± standard deviation are shown.

| Atlas (# of ROIs) | Accuracy (%) | AUC |
| --- | --- | --- |
| FS86 (86) | 72.95 ± 3.33 | 0.7864 ± 0.0388 |
| TT (94) | 72.95 ± 4.89 | 0.7831 ± 0.0438 |
| HO (110) | 74.23 ± 5.25 | 0.8116 ± 0.0334 |
| AAL (116) | 74.10 ± 2.08 | 0.7924 ± 0.0278 |
| EZ (116) | 74.49 ± 4.81 | 0.8037 ± 0.0401 |
| CC200 (200) | 77.56 ± 4.69 | 0.8511 ± 0.0355 |
| CC400 (392) | 81.15 ± 3.53 | 0.8655 ± 0.0325 |
| **Ensemble Model** | **81.15 ± 3.39** | **0.8634 ± 0.0333** |

**Table 2:**
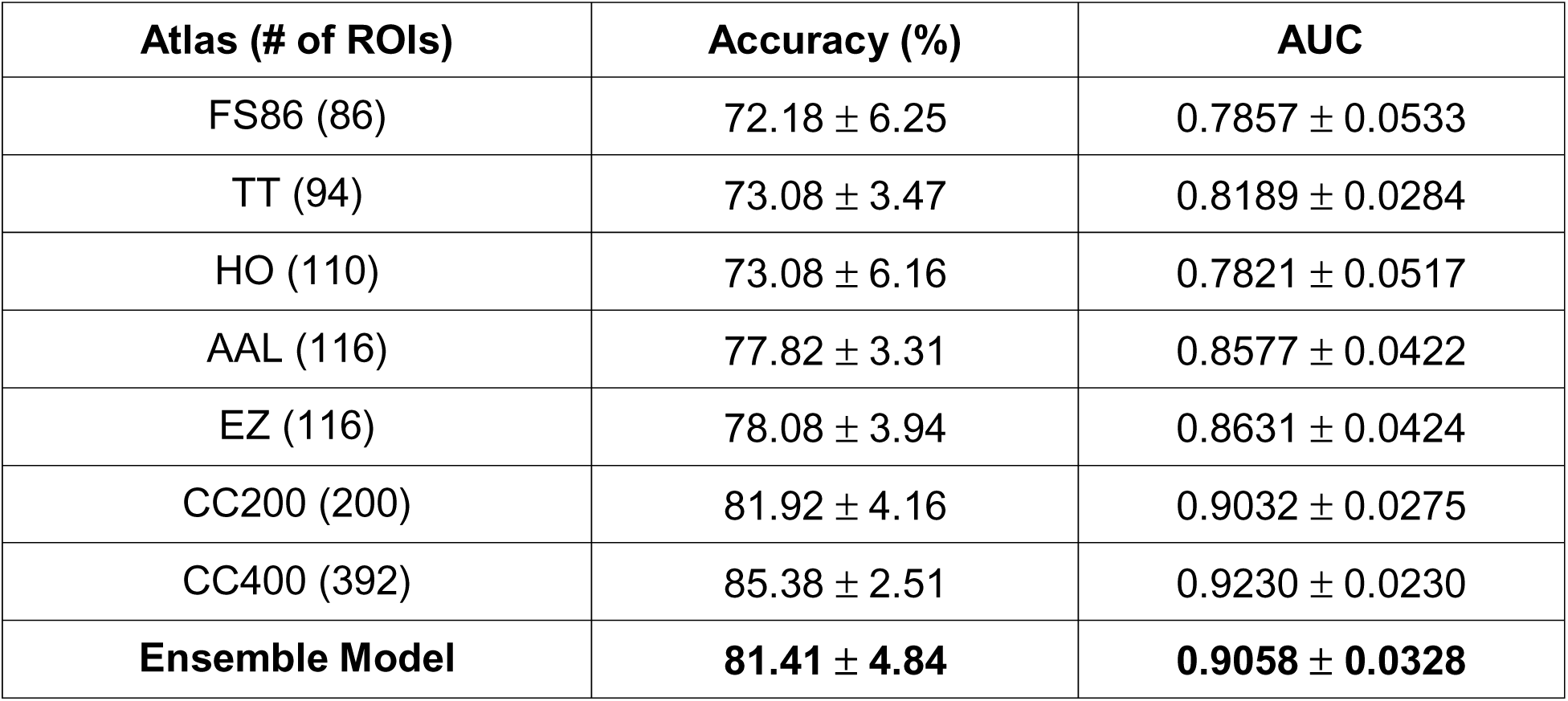
Results from Grey Matter Volume Matched Dataset (n=390). Using ROI volume to predict sex. Ten permutations completed for each atlas. Mean accuracy (balanced) ± standard deviation and mean AUC ± standard deviation are shown.

Models trained on HE from different atlases identified similar cortical and subcortical regions as being important features for sex classification. A feature importance map for HE-based classification for the CC400 atlas is shown in Figure 4a, and for all other atlases in Figure S4. Regions exhibiting the strongest feature importance were found in the cerebellum, amygdala, frontal cortex, and occipital cortex. Models trained on volume measures derived from different atlases also identified similar cortical and subcortical regions as being important features for sex classification. A feature importance map for volume-based classification for the CC400 atlas is shown in Figure 4b, and for all other atlases in S5. Regions exhibiting the strongest feature importance were found in the cerebellum, cingulate cortex, and temporal cortex. Feature importance for HE-based models and volume-based models were not correlated for any of the atlases (p-corrected > 0.05).

**Figure 4.**
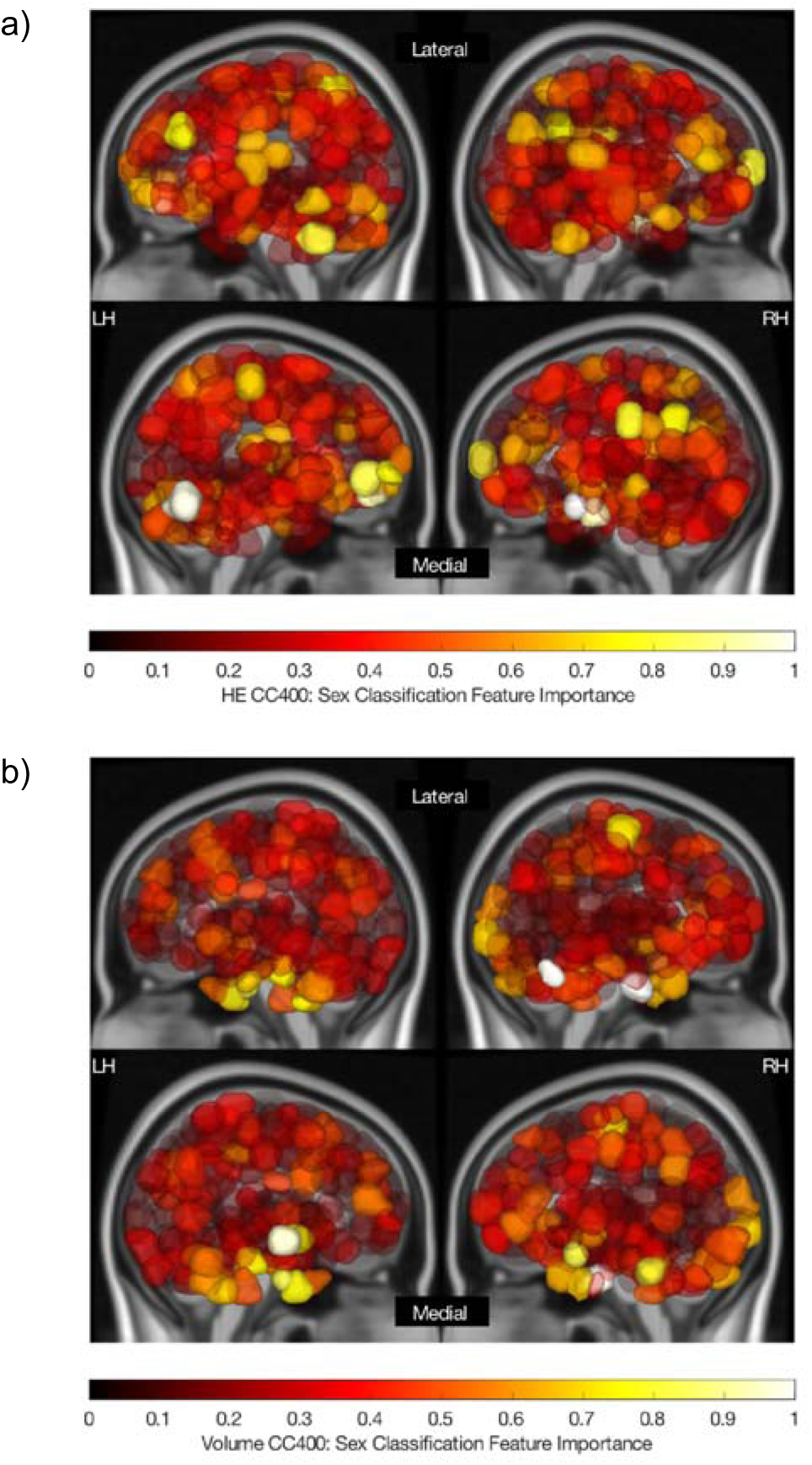
Feature importance map for a linear SVM classifier used to predict sex using HE (a) and volume (b) computed on the CC400 atlas. Lateral (top) and medial (bottom) sides of the left (LH) and right (RH) hemispheres are shown for both a and b. The absolute value of feature weights obtained from the linear SVM were scaled to generate feature importance values as plotted per the colour scale. Values closer to 1 indicate greater importance in the overall classification.

Results from the atlas-specific ensemble models combining the predicting probabilities from the HE- and volume-based classification models are shown in Table 3. The CC400 atlas’ ensemble model performed with a mean accuracy of 87.3% (SD=3.2%) and a mean AUC of 0.94 (SD=0.015). The atlas-specific ensemble models significantly outperformed the HE-based models in terms of accuracy (p<0.05) and AUC (p<0.01). While these atlas-specific ensemble models also had a higher mean accuracy and AUC than the volume-based models, this difference was not significant (p>0.05).

**Table 3:** Results from Grey Matter Volume Matched Dataset (n=390). Using atlas-specific ensemble models to combine prediction probabilities from the HE- and volume-based classification models for each test subject’s sex. Ten permutations completed for each atlas. Mean accuracy (balanced) ± standard deviation and mean AUC ± standard deviation are shown.

| **Atlas (# of ROIs)** | **Accuracy (%)** | **AUC** |
| --- | --- | --- |
| FS86 (86) | 75.77 ± 6.34 | 0.8610 ± 0.0376 |
| TT (94) | 78.59 ± 2.57 | 0.8698 ± 0.0257 |
| HO (110) | 77.69 ± 4.70 | 0.8581 ± 0.0446 |
| AAL (116) | 82.31 ± 4.28 | 0.8931 ± 0.0317 |
| EZ (116) | 82.56 ± 4.34 | 0.8986 ± 0.0314 |
| CC200 (200) | 86.15 ± 4.24 | 0.9327 ± 0.0261 |
| CC400 (392) | 87.31 ± 3.22 | 0.9364 ± 0.0151 |
| **Ensemble Model** | **86.67 ± 4.22** | **0.9330 ± 0.0274** |

Models trained on HE from the CC400 atlas for entire dataset (n=1003) had a mean balanced accuracy of 85.4% (SD=2.1%) and mean AUC of 0.94 (SD=0.012). Models trained on HE from the CC400 atlas for 10 randomly selected (not volume-matched) sample-size-matched subsets (n=390) performed with a mean balanced accuracy of 84.0% (SD=1.7%) and a mean AUC of 0.92 (SD=0.012). A comparison of the balanced accuracy and AUC distributions obtained from sex classification models based on HE on the grey-matter-volume-matched subset, random sample-size-matched subset, and the entire dataset are shown in Figures 5a and 5b. Mean balanced accuracies achieved by models trained with the grey-matter-volume-matched subset were significantly worse than models trained with the entire dataset (p<0.01). Mean balanced accuracies of models trained with sample-size-matched subsets were not significantly different from models trained with the grey-matter-volume-matched subset (p>0.05) or models trained with the entire dataset (p>0.05). Mean AUCs achieved by models trained with the grey-matter-volume-matched subset were significantly worse than models trained with the sample-size-matched subsets (p<0.001) and models trained with the entire dataset (p<0.001). Mean AUCs of models trained on the sample-size-matched subsets and the entire dataset were not significantly different (p>0.05).

**Figure 5:**
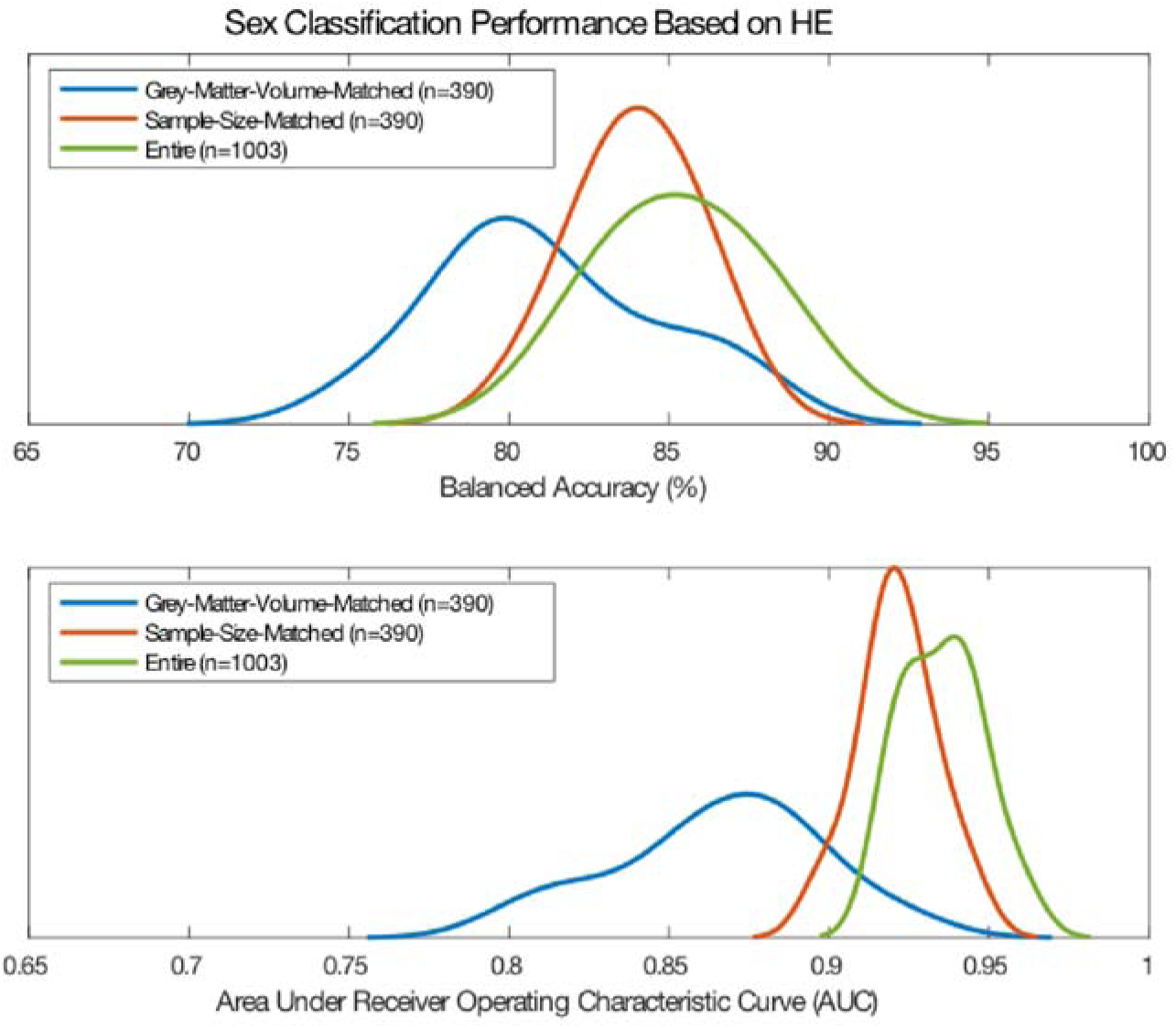
Balanced accuracy (a) and AUC (b) distributions obtained from sex classification models based on HE on the grey-matter-volume-matched subset, randomly sample-size-matched-subset, and the entire dataset.

## Discussion

Our findings reveal sex differences in long-range temporal dependence in healthy young adults, with males exhibiting more persistent long-range temporal dependence (higher HE) than females in the temporal, parietal, and occipital cortices. We performed sex classification based on regional temporal dependence using a linear SVM and achieved mean accuracy as high as 81.1% and mean AUC as high as 0.86 across all atlases. Additionally, we found that even when males and females are strictly matched on total grey matter volume, significant sex differences persist in regional volume across many cortical and subcortical regions. In fact, sex classification based on regional volume using a linear SVM and achieved mean accuracy as high as 85.3% and mean AUC as high as 0.92.

Previous work has found that temporal dependence of brain activation time series varies with states of consciousness [12] and exhibits age and sex effects [7], but the cognitive and/or behavioral implications of temporal dependence have not yet been explicitly studied. Additionally, biological substrates driving HE are not well understood. Although sex differences at cellular and molecular levels of the nervous system have been observed [13], we do not know whether it is gene expression, neuronal signalling, glial activity, anatomical structure, regional blood flow, hormone effects or other factors that are driving them. It is even less clear how these factors may contribute to sex differences in temporal dependence, as measured by HE, within the resting-state time series. As a result, it is difficult to interpret what these results may represent in a biological or behavioural domain.

While no previous studies, to our knowledge, have attempted sex classification using HE, studies have used ROI-based resting-state functional connectivity measures for the classification using this same dataset [7, 8]. Those studies reported maximum test accuracies of 86.6% (AUC of 0.93) using whole-brain functional connectivity [7] and 72.6% using region-specific functional connectivity [8], which are comparable to the results reported here. This shows that while HE and functional connectivity represent different information about activation patterns in the brain, they share the ability to make accurate predictions about sex. HE is inherently lower in dimensionality, thus avoiding the curse of dimensionality, and can be mapped to a single specific region in the brain, making it potentially easier to interpret than functional connectivity that represents pairs of regions.

One key difference between our study and previous ones is that we strictly control for volume differences between the sexes. This is a major factor to consider when studying sex differences as there is inherent bias introduced by volumetric differences between males and females [13, 14]. In an initial analysis performed on the entire HCP dataset (n=1003), we found that 725 regions (out of 1114 across seven atlases) exhibited significant sex differences in HE. The regions identified in this analysis matched those previously observed [3]. However, upon repeating the analysis on the grey-matter volume matched subset (n=390) to eliminate the effects of volumetric differences, we found that only a subset of the originally identified regions were significantly different between males and females. This suggests that previous results [3] may be confounded by volume differences and these differences must be accounted for when analysing sex differences.

In terms of the influence of volumetric differences on sex classification, a previous study using functional connectivity to classify sex differences reported a decrease in performance in a grey-matter-volume-matched subset compared to their entire dataset [8]. However, they attributed the decrease in performance to the overall decrease in sample size. Our results show that the mean AUC for the sex classification task was significantly lower for models trained with the grey-matter-volume-matched subset compared to both the entire dataset and the randomly selected (not volume-matched) subsets of the same size. This suggests that, in our case, the decrease in AUC cannot be attributed to the decrease in sample size; rather, volume-matching reduced the classification performance of our fMRI-only based biomarker. Future studies that investigate sex differences in functional activation patterns must be aware that results may be inflated if they are not strictly volume-matching their female and male populations.

Studies often tend to use global volume matching strategies to reduce sex effects on volume. We show here that even after matching subjects on total grey matter volume, significant volumetric sex differences persist in many cortical and subcortical regions. Males had significantly larger volumes in the cerebellum, hippocampus, parahippocampus, thalamus, caudate, and amygdala while females had significantly larger volumes in the cingulate, precuneus, frontal cortex, and parietal cortex. This demonstrates that sex differences in regional brain volume exist even after matching subjects for grey matter volume. Finally, atlas-specific ensemble models that combined the prediction probabilities from the HE- and volume-based models had significantly better performance than HE based models and trends for better performance than volume based models. This suggests that HE and volume measures are capturing distinct information that distinguishes males and females.

Male and female brains are similar in many respects but very different in others [13]. Understanding the biology of male and female brain functionality in healthy individuals is crucial to elucidating mechanisms and determining more effective interventions for many neurodevelopmental, neurological, and psychiatric illnesses, which exhibit sex differences in prevalence, age of onset, and symptomatology [15]. Both genetic and hormonal influences play a role in brain development, but any sex difference in neural structure may also be shaped through experience, practice, and neural plasticity [13]. In this study, sex differences were observed in the long-range temporal correlation of resting-state time series. However, the relationship between genetic information, hormonal measures, and HE was not evaluated in this study. In the absence of proof of genetic or hormonal influence underlying these observed sex differences in HE, it is important to note that any existing sex difference may have been shaped through experience, practice, or neural plasticity [16].

### Limitations

Machine learning problems based on neuroimaging data are prone to the curse of dimensionality. Voxel-wise data are on the order of hundreds of thousands of features, and even when examining ROI-based data, there can be several hundred features. To avoid the curse of dimensionality and remove noise from the data, we used parcellations of the brain to generate a single measure per ROI for each subject. This resulted in the dimensionality being reduced from hundreds of thousands to between 86 and 392, depending on the atlas. However, by drastically reducing the overall dimensionality, information may have been lost and biases may have been introduced that limit the overall classification performance. In an attempt to mitigate possible biases introduced by atlas selection, the findings described in this paper were evaluated over seven different parcellation schemes. However, to further reduce possible bias introduced by the use of atlases, a voxel-wise analysis of sex differences could be examined in future work. Finally, a single scaling parameter such as the HE is limited in its ability to characterise temporal features due to the complex nature of fMRI time series [17]. An alternative to the HE is to use a whole set of fractal Hölder exponents instead which can also account for local intensity fluctuations [17].

Sex differences in functional brain activity may be due to an underlying hormonal effect. While various studies have shown that resting-state functional connectivity fluctuates across the menstrual cycle in women [18–20], the effect of hormones on the HE has not yet been studied. Hormonal measures which would allow the study of the relationship between females’ menstrual cycles and regional HE were not available in the dataset used for this work. Future datasets should aim to collect hormonal levels such that a thorough investigation on the effect of menstrual cycle on HE can be examined. It is also important to note that hormonal fluctuations in regional HE may result in females exhibiting increased variability in those regions, as conjectured in [8]. Consequently, this could influence classification performance such that classification is more accurate in women during certain points of their cycles than others.

Studies examining sex differences often do not consider an individual’s gender identity and fluidity. This study only used information about each subject’s self-reported sex in the absence of gender identification. Males and females are exposed to different expected gender roles and a lifetime of gender-differentiated experience could be the underlying cause to sex differences in neuroimaging biomarkers [16]. These social factors may be partially responsible for the regional group differences in temporal dependence and overall classification performance reported in this study. Future datasets should aim to collect data about gender identity and fluidity for subjects and studies should incorporate this information into their work.

A recent study used resting state functional connectivity data from HCP dataset to make predictions about behavioural function [21]. The study identified that using functional connectivity with global signal regression improves prediction of a wide range of behavioural phenotypes compared to functional connectivity without global signal regression. It is therefore important to acknowledge that preprocessing steps used in the HCP data as well as in computation of HE for this study may have an influence on the overall results obtained.

Lastly, this study only used data from the HCP dataset. Time series obtained from fMRI can be sensitive to scan parameters and pre-processing pipelines. Although test hold-out sets were used exclusively for evaluating the models generated for sex classification, the overall results may not be entirely generalisable to other datasets.

## Conclusion

Understanding sex-specific brain differences in healthy individuals is a critical first step towards understanding sex-dependent variation in neurological, developmental, and psychiatric disorders and, possibly, the use of this information to develop personalised interventions. In this study, we observe that males exhibit higher temporal dependence of resting-state time series than females in the temporal, parietal, and occipital lobes. Furthermore, using ROI-based information about temporal dependence, we were successfully able to classify males and females using a linear SVM algorithm with a maximum mean accuracy of 81.1% and mean AUC of 0.86. We also identify that regional volume differences between males and females persist even after matching for cortical and subcortical grey-matter volume. Regional volume can also be used to successfully classify males and females with a maximum mean accuracy of 85.3% and mean AUC of 0.92. Finally, we demonstrate that matching male and female groups for grey matter volume can decrease sex-classification accuracy of functionally-derived biomarkers, an issue that must be a carefully considered in future studies. Additional research is needed to understand the biological substrates underlying the variations in temporal dependence and comprehend the behavioral or cognitive implications of these sex differences.

## Materials and Methods

An outline of our overall workflow can be found in S1. All codes used for data analysis are available on GitHub (https://github.com/elvisha/HurstSexDifferences).

### Dataset

Publicly available high-resolution magnetic resonance imaging data from the Human Connectome Project – Young Adult S1200 release [9] were used in this study. We examined time series from 1003 volume-matched healthy adults having four complete resting-state fMRI runs (1200 volumes each). The subjects had a mean age of 28.71 years (range: 22 – 37 years, SD=3.71, median =29) and included 469 males (46.8%). In order to avoid biases introduced by volumetric differences between males and females, a subset of 390 volume-matched subjects (190 non-overlapping male-female pairs) were identified and all analyses for this study were performed solely on that subset. The mean age of the 390 subjects was 28.6 years (range 22-37 years, SD=3.7, median=29).

Although the term “gender” is used in the HCP data dictionary, we use the term “sex” in this paper because the database collected subject self-reported biological sex as opposed to gender identification. Genetic information was not used to verify the self-reported biological sex.

### Preprocessing of fMRI Data

HCP MR imaging data were acquired on a Siemens Skyra 3T scanner at Washington University in St. Louis. Each subject underwent four gradient-echo EPI rfMRI runs (TR = 720 ms, TE = 33.1 ms, 2.0 mm isotropic voxels, FoV = 208 × 180 mm^2^, flip angle = 52 degrees, matrix = 104 × 90, 72 slices) of about 15 min each over two sessions: two runs in the first sessions (REST1_LR and REST1_RL) and two runs in the second session (REST2_LR and REST2_RL). The data consisted of 1200 volumes for each run for a total of 4800 volumes for each subject over the four runs. Each run of each subject’s rfMRI was preprocessed by the HCP consortium [10]. The data was minimally preprocessed [22] and had artefacts removed using ICA+FIX [23, 24].

### Motion

Framewise displacement [25] (FD) was computed for each subject. Sex differences in framewise displacement for the specific windows used to calculate HE were evaluated using a two-sample t-test. There were no sex differences identified in motion.

### Parcellations

As part of the HCP preprocessing pipeline [22], FreeSurfer’s recon-all pipeline [26–32] was optimised for the high-resolution HCP data. Structural T1-weighted and T2-weighted images (0.7mm isotropic) were used with the optimised HCP FreeSurfer pipeline to automatically segment subcortical structures [29] and cortical sulci and gyri [33]. These segmentations were used to reduce the fMRI data of the whole brain into cortical and subcortical grey matter regions of interest (ROI). For ROI-based analyses, the following 7 different parcellations were used, where R denotes the number of ROIs: Automated Anatomical Labeling [34] (AAL, R=116), Craddock 200 [35] (CC200, R=200), Craddock 400 [35] (CC400, R=392), Eickhoff-Zilles [36] (EZ, R=116), Harvard-Oxford [37, 38] (HO, R=110), Talairach and Tournoux [39] (TT, R=94), as well as a FreeSurfer aparc+aseg parcellation (FS86, R=86) in which cortical grey matter was divided into 34 regions per hemisphere as per the Desikan-Killiany atlas [33], and subcortical grey matter was divided into 9 regions per hemisphere.

### Hurst Exponent

Detrended fluctuation analysis (DFA) was used to study the temporal correlations of BOLD fluctuations and calculate the Hurst Exponent (HE) as previously described [1] and summarised below.

1. Given *x_t_*, series of *T* measurements, 1 ≤ *t* ≤ *T*, subtract mean of signal, < *x* >, and compute cumulative sum, 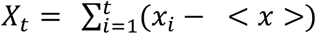.
2. Divide signal into non-overlapping windows of length *L*, with each segment labeled as 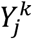, with *j* indexing time within segment (1 ≤ *j* ≤ *L*) and *k* indexing segment number (1 ≤ *k* ≤ *T*⁄*L*).
3. For each segment, fit a linear function using least squares to determine slope, *a^k^* and intercept, *b^k^*. Subtract best fitting linear trend and compute fluctuation from mean of resulting signal, 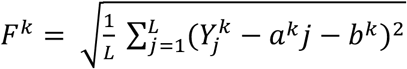.
4. Average *F^k^* for all segments at temporal scale *L* to yield fluctuation function, *F*(*L*).
5. Repeat 2-4 using different window lengths, *L* = [10, 15, 25, 30].
6. The slope of *F*(*L*) in logarithmic scale vs. *L* is the HE.

For every subject, DFA was applied to 32 segments (8 segments per scan) of 150 volumes each of the BOLD time series of every voxel. The HE computed over the 32 segments were averaged together to generate a single mean HE for each voxel. The HE was averaged over voxels within each ROI to generate a single HE value per ROI.

### Volume

To compare ROI size differences, subject-specific ROI volumes were computed by counting voxels in each subject’s native 0.7mm anatomical space. The FreeSurfer-derived FS86 atlas was already defined in this native space. The remaining atlases were defined in 2mm MNI152 space, and were first transformed from MNI152 to each subject’s native space using the standard2acpc_dc nonlinear warp provided by HCP and nearest-neighbour interpolation.

To avoid biases introduced by volumetric sex differences, 390 subjects (190 non-overlapping male-female pairs) were selected such that each pair had a matched total grey matter volume (cortical and subcortical) with a percent difference less than or equal to 1%. The final volume-matched sample did not differ in total grey matter volume (p>0.05) between males and females.

### Sex Differences in HE and Volume

For each atlas, sex differences in HE across the ROIs were evaluated. For each ROI, the difference in means between males and females was analysed using a two-tailed two-sample *t*-test. This was repeated with all atlases. P-values were adjusted for multiple comparisons across all atlases using the Benjamini-Hochberg procedure [11] to decrease the false discovery rate. This same process was repeated for volumetric sex differences.

### Sex Classification

In order to classify subjects in the grey-matter-volume-matched subset into males and females using ROI-based HE or regional volume from each atlas, a linear Support Vector Machines (SVM) approach was implemented [40]. SVMs [41] are supervised learning models which can be used for classification tasks. Given labeled training data with *n* features, the algorithm outputs an optimal hyperplane which can be used to separate the classes with a maximal margin. If the data are linearly separable, linear SVMs can be used to find this optimal hyperplane. Linear ridge, SVM with radial basis function kernel, random forest, and neural network classifiers were also evaluated in this work. However, the performance metrics obtained from those classifiers were worse than those obtained for linear SVM so the results for this paper focus on linear SVM.

Subjects were separated into stratified train (80%) and test (20%) subsets and the same train and test subsets were used across all atlases so that results could be directly compared with one another. For each atlas, hyperparameter tuning of penalty parameter C was conducted using nested grid search cross validation in which both the inner and outer folds were randomly split into 5 stratified groups. A single best model and the hyperparameter C corresponding to it were identified. A coarse grid search followed by a finer grid search across the same hyperparameter space was conducted for all atlases. This nested grid search cross validation was repeated 100 times to generate 100 separate values for C. The final model was created by averaging the hyperparameter C obtained across 100 iterations of the finer grid search. This final model was evaluated on the test subset and the corresponding accuracy, area under the receiver operating characteristics curve (AUC), and feature importance are reported. This was repeated using the same 10 permutations of randomised training and testing splits for all atlases to get a distribution of overall performance metrics for both HE- and volume-based classification. The feature weights vector obtained from the linear SVM classifier for each atlas was averaged across the permutations and the absolute value of the average scaled such that feature importance for each ROI falls between zero and one, and it is easier to interpret across atlases. An ensemble model that combined prediction probability for each test subject’s sex across all atlases [42] was also implemented for both HE- and volume-based sex classification separately. Atlas-specific ensemble models that combined prediction probability from the HE- and volume-based classification for each test subject’s sex was also implemented. Lastly, a final ensemble model that combined prediction probability from both the HE- and volume-based classification for each test subject’s sex across all atlases. Differences in the classification AUC and accuracy between the HE-based classification, volume-based classification, and ensemble model classification based on both HE and volume were evaluated using a one-way analysis of variance (ANOVA).

Sex classification based on HE was also performed for the entire dataset and randomly sample-size-matched subsets using the CC400 atlas. Ten permutations of sex classification, as described above, were completed using stratified splits of the entire dataset (n=1003, 46.8% male) to determine the overall effect of grey matter volume matching on the performance metrics. Additional permutations of sex classification using randomly generated subsets were also performed to determine how an overall decrease in sample size (unrelated to grey matter volume matching) may affect the results. Ten distinct subsets (n=390) were randomly generated by selecting 195 males and 195 females from the entire dataset. For each of the ten distinct subsets, ten permutations of sex classification, as described above, were completed. Balanced accuracy and AUC distributions were generated by averaging the results across the subsets. Finally, differences in the classification AUC and balanced accuracy between the grey-matter-volume-matched subset, randomly sample-size-matched subset, and entire dataset were evaluated using a one-way ANOVA.

## Acknowledgements

Data were provided by the Human Connectome Project, WU-Minn Consortium (Principal Investigators: David Van Essen and Kamil Ugurbil; 1U54MH091657) funded by the 16 NIH Institutes and Centers that support the NIH Blueprint for Neuroscience Research; and by the McDonnell Center for Systems Neuroscience at Washington University. This work was supported by the following grants: R21 NS104634-01 (AK) and R01 NS102646-01A1 (AK). The authors would also like to acknowledge Clive Aaron D’Souza and Shashank Pathak from Cornell University for their contributions to an earlier version of this work.

