## Supplemental Figures for "Sex classification using long-range temporal dependence of resting-state functional MRI time series"

S1. Overall workflow of analysis. Functional MRI images (1) from the Human Connectome Project were used to extracted voxel-wise time series (2). Voxel-wise Hurst exponents were computed (3). Parcellations of the voxels were generated for seven different atlases (4) and Hurst exponents were averaged for all voxels within a given ROI to generate ROI-based Hurst exponents (5). Sex differences in regional Hurst exponent were analysed using a Student’s t-test (6). Prediction of sex was performed using a linear SVM classifier (7). Nested cross validation was used to optimize hyperparameters and a final model was fitted to the train data and evaluated on the test data.

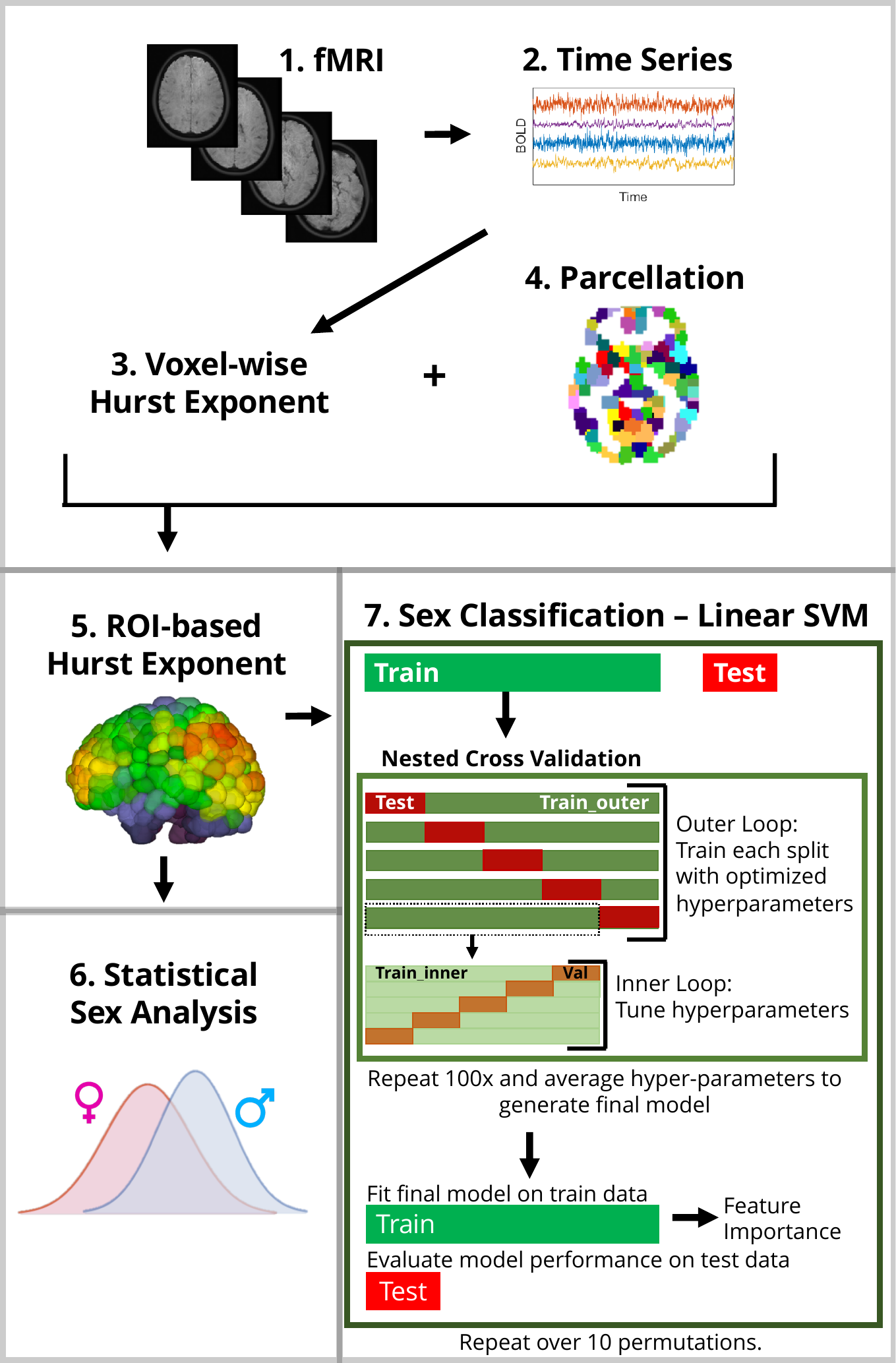

S2. Region-wise sex differences in HE in grey matter computed on the AAL (a), CC200 (b), EZ (c), FS86 (d), HO (e), and TT (f) atlases. Lateral (top) and medial (bottom) sides of the left (LH) and right (RH) hemispheres are shown for each atlas. Regional t-statistics are shown as per the colour scale for all significantly different (p-corrected < 0.05) areas. Non-significant areas are shown in grey. A positive t-statistic indicates that males have a higher mean HE value in that region than females.

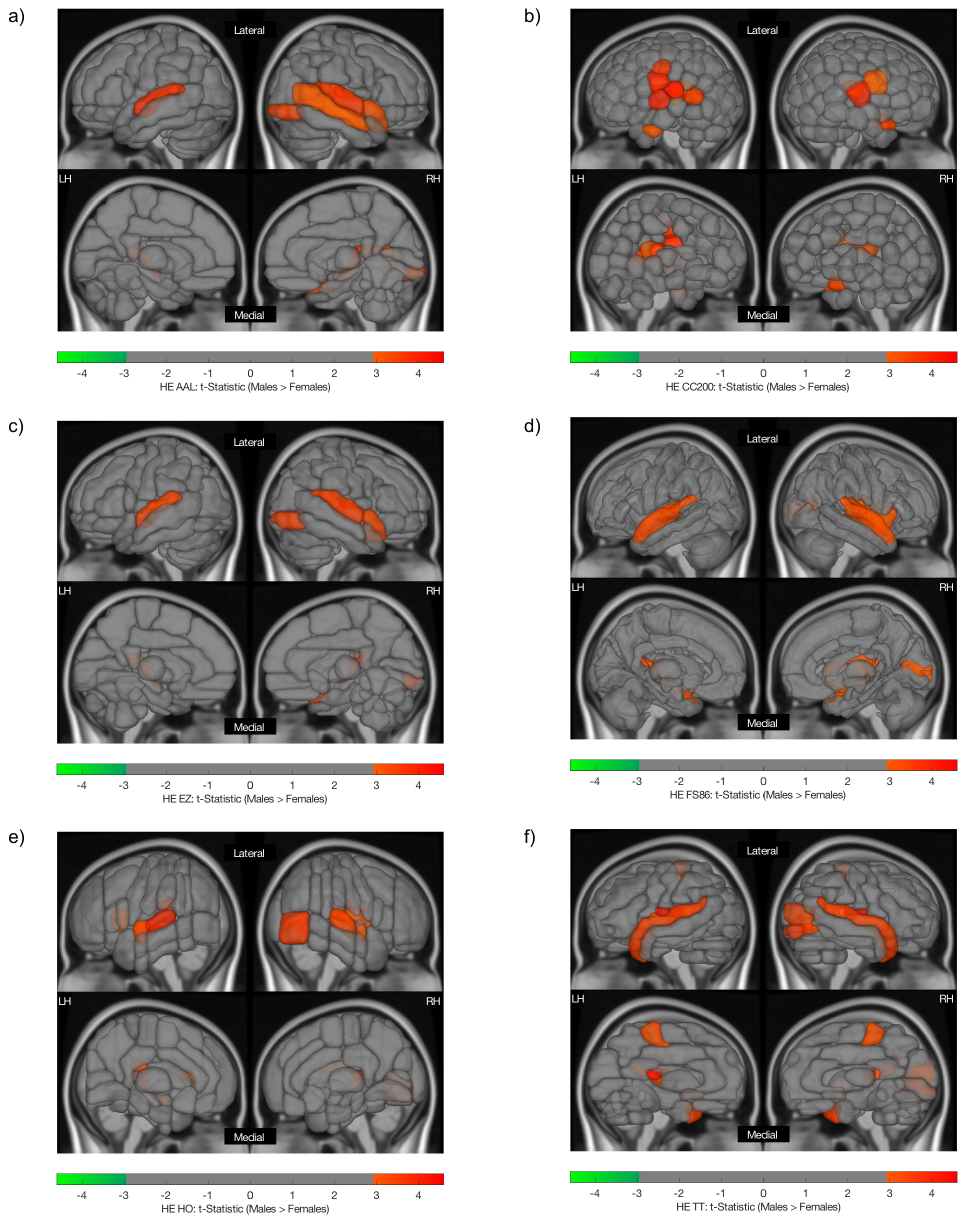

S3. Region-wise sex differences in volume computed on the AAL (a), CC200 (b), EZ (c), FS86 (d), HO (e), and TT (f) atlases. Lateral (top) and medial (bottom) sides of the left (LH) and right (RH) hemispheres are shown for each atlas. Regional t-statistics are shown as per the colour scale for all significantly different (p-corrected < 0.05) areas. Non-significant areas are shown in grey. A positive t-statistic indicates that males have a higher mean regional volume than females.

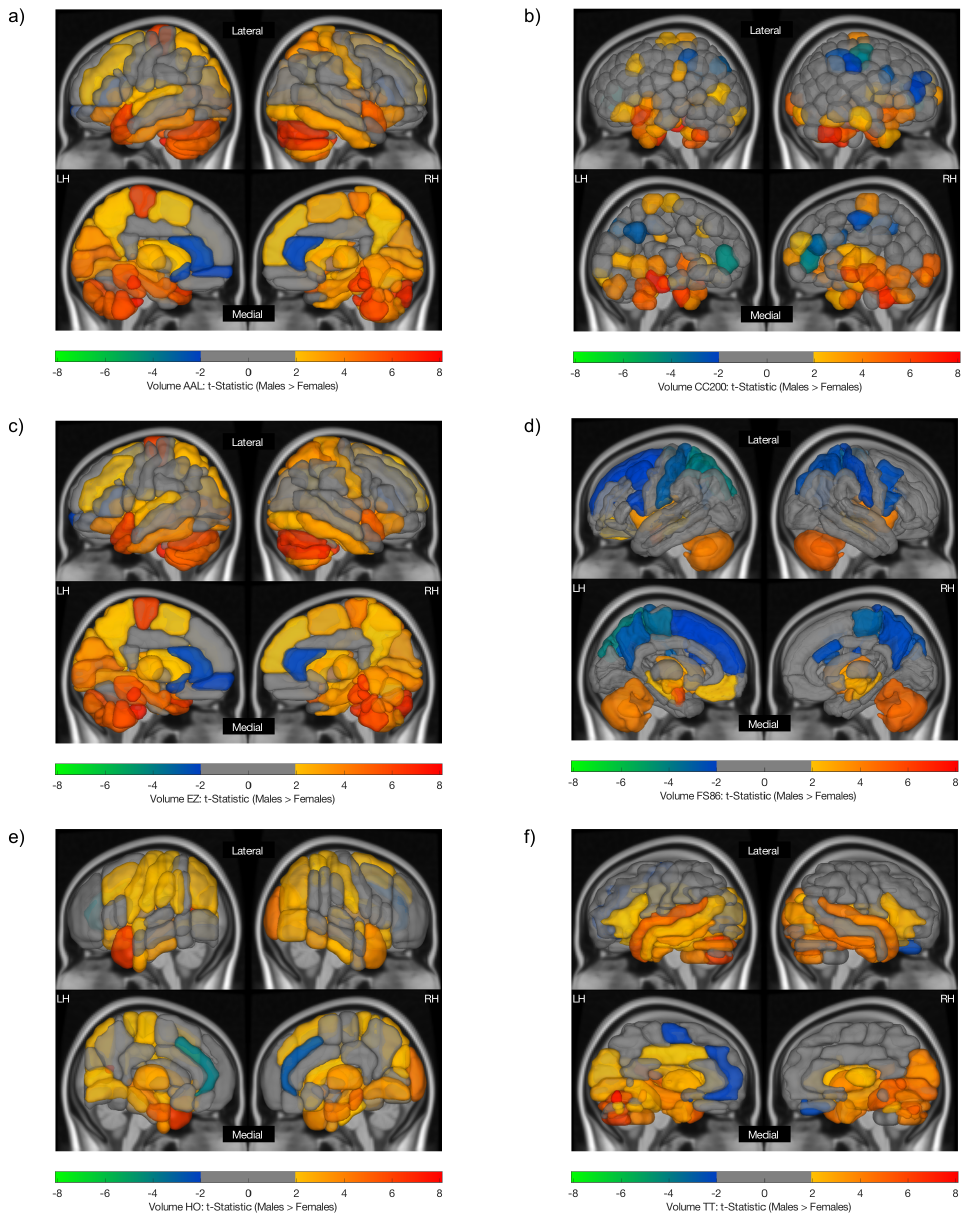

S4. Feature importance map for a linear SVM classifier used to predict sex using HE computed on the AAL (a), CC200 (b), EZ (c), FS86 (d), HO (e), and TT (f) atlases. Lateral (top) and medial (bottom) sides of the left (LH) and right (RH) hemispheres are shown for each atlas. The absolute value of feature weights obtained from the linear SVM were scaled to generate feature importance values as plotted per the colour scale. Values closer to 1 indicate greater importance in the overall classification.

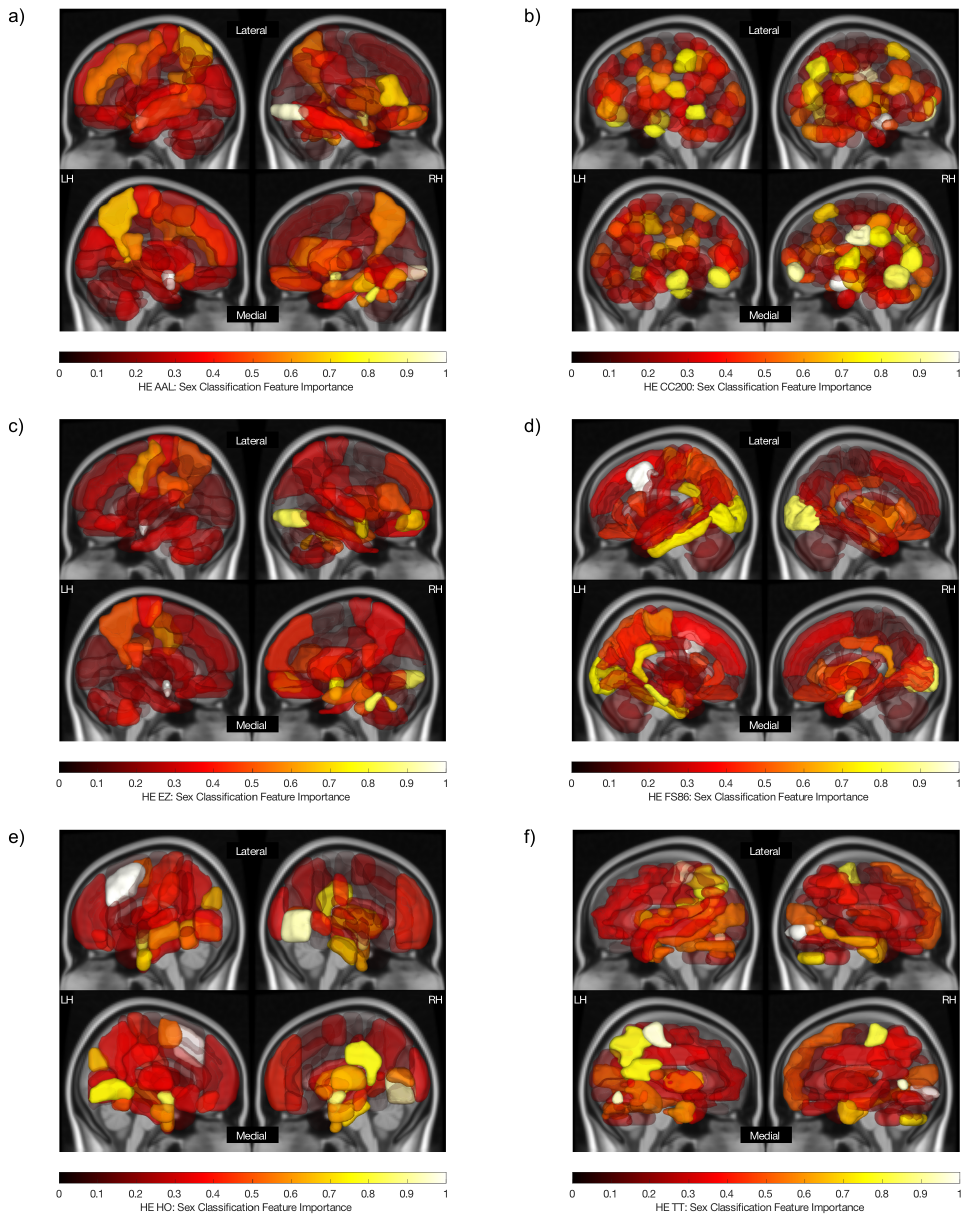

S5. Feature importance map for a linear SVM classifier used to predict sex using volume computed on the AAL (a), CC200 (b), EZ (c), FS86 (d), HO (e), and TT (f) atlases. Lateral (top) and medial (bottom) sides of the left (LH) and right (RH) hemispheres are shown for each atlas. The absolute value of feature weights obtained from the linear SVM were scaled to generate feature importance values as plotted per the colour scale. Values closer to 1 indicate greater importance in the overall classification.

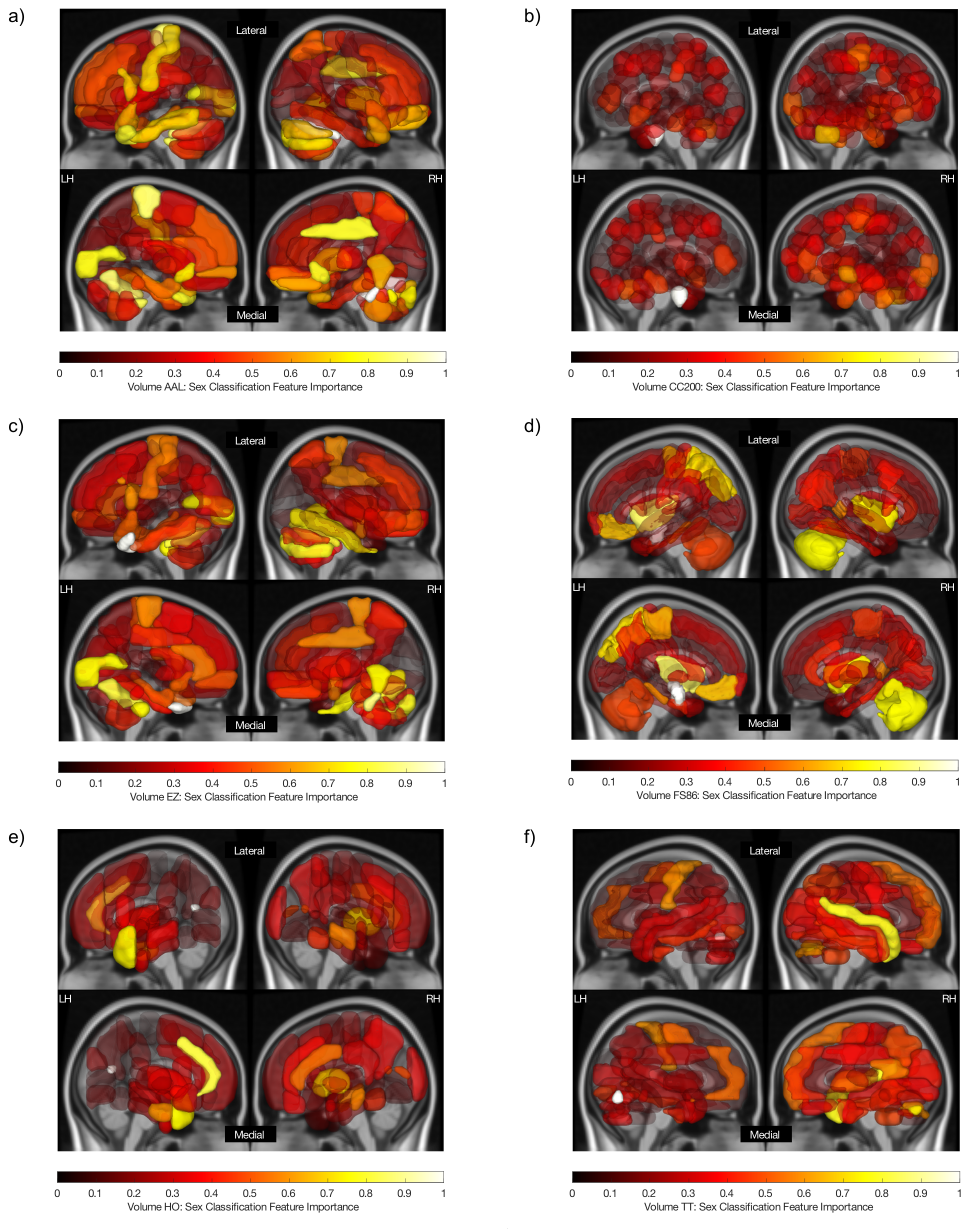
